# Response of mate harm to sex-separated gene pools

**DOI:** 10.1101/2025.09.29.678620

**Authors:** Chloe Melo-Gavin, Mathew P. Lindsay, Aneil F. Agrawal

**Affiliations:** Department of Ecology and Evolutionary Biology, University of Toronto, Toronto, Ontario, M5S 3B2 Canada

**Keywords:** sexual antagonism, sexual differential selection, mate harm

## Abstract

Competition among males to fertilize females can generate sex differences in selection. At the genetic level, alternative alleles can be favoured in each sex (intralocus sexual conflict, IASC). At the phenotypic level, males can evolve traits that are harmful to the females with whom they interact. Here we used experimental evolution in *Drosophila melanogaster* to examine how relaxing IASC affected the evolution of mate harm. Genetically variable Chromosome 2s evolved in two separate pools within each population. One pool experienced patrilinear inheritance (segregated like a Y-chromosome) and male-limited selection, while the alternative pool segregated like an X-chromosome and experienced female-biased selection. We measured female fitness when exposed to males carrying either chromosome type at either normal (continuous) or reduced (periodic) male exposure, over three time periods. Males carrying either type of chromosome were harmful to females, but males carrying a male-limited chromosome displayed increased harmfulness, suggesting ongoing selection for mate harm that is constrained under normal inheritance. The magnitude and direction of the effect of male interaction on female fitness was highly sensitive to time of measurement, and we observed a brief period where increased interaction with males was beneficial to females.

## Introduction

Divergent selective pressures between the sexes can lead to opposing strategies for maximizing fitness. At the genetic level, alleles at shared loci may have conflicting fitness effects in males and females (Parker, 1979). This sexual antagonism, termed intralocus sexual conflict (IASC), can then prevent the sexes from fixing their preferred allele due to its effect in the other sex (Rice and Chippindale, 2001) and occurs in a wide range of phenotypes (Long and Rice, 2007; Abbott et al., 2010; Okada et al., 2021; Arnqvist and Rowe, 2005; Mainguy et al., 2009). Additionally, conflict over reproductive interactions can select for traits that increase the fitness of one sex even if it damages the fitness of the other (e.g. a male trait that increases his mating success even though it reduces the survival of females with whom he interacts). These mate harming traits have been documented for a diverse array of phenotypes from aggressive courtships (Fielding and Knisley, 1995) to traumatic matings (Le Boeuf and Mesnick, 1991; Kamimura, 2007) and toxic seminal fluid (Chapman et al., 1995; Partridge et al., 1986; Sirot et al., 2014).

Frequently, separate sex-limited traits mediate male-female interactions (e.g., grasping hooks in male water-striders and anti-grasping armaments in females (Arnqvist and Rowe, 2002). It is often assumed that different loci underly these separate traits and, as such, conflict occurring at the level of inter-individual interactions has been termed “interlocus sexual conflict” to distinguish it from IASC (Bonduriansky and Chenoweth, 2009). Perhaps because of these seemingly mutually exclusive labels, the interaction of IASC (conflict over shared genetics) and mate harm (inter-individual conflict) has been relatively understudied, though others have pointed out they may be genetically related (Pennel et al., 2016; Romero-Haro et al., 2023; Jiang et al., 2010) Even if a trait involved in mate harm is sex-limited (e.g., aggressive courtship behaviour by males), the alleles affecting that trait may have pleiotropic effects on other traits, including those expressed in the other sex. Indeed, mate harm ability is mediated, at least in part, via traits present in both sexes (e.g., body size, locomotor activity, etc.; Pitnick and García– González, 2002; Nandy et al., 2013). Because the selection on the trait (e.g., courtship persistence) that causes harm to the members of the other sex is usually sex-specific, this will typically result in the underlying loci being differentially selected between the sexes and thereby potentially subject to IASC. For example, locomotor activity has opposing effects on male and female fitness in *D. melanogaster* (Long et al., 2007). The positive selection in males exists because locomotion facilitates their mate search and courtship of females. Thus, an allele that increases locomotion in both sexes (and subject to IASC) would also increase male-female interaction and mate harm. For simplicity, we have discussed this connection between IASC and mate harm with respect to traits that cause harm, but the same logic applies for traits involved in the resistance to harm.

The connection between mate harm and IASC is important for both topics. As stated above, the selection that causes the evolution of mate harming traits (or traits for resisting such harm) is often sex-specific at the trait level and may cause IASC at the gene level. Conversely, IASC can affect the evolution of mate harm. When Pennell et al. (2016) modelled the co-evolutionary dynamics of mate harm, they found that incorporating pleiotropic effects of mate harming traits in both sexes (i.e., allowing for IASC) altered classic predictions of interlocus sexual conflict, and, in some cases, halted or reversed the evolution of mate harm.

While it is plausible that the connection between mate harm and IASC may be common, empirical evidence is limited. Romero-Haro et al. (2023) found that artificial selection in female quail for female-limited traits such as egg investment, can produce correlated changes in male traits affecting mate harm such as male ejaculate toxicity, demonstrating that even sex-limited mate harming traits can be affected by selection in the other sex. However, that study did not aim to evaluate whether this reflects a connection between mate harm and IASC *per se*.

Experimental evolution offers a means to test for this connection between IASC and mate harm. Because IASC can prevent the sexes from fixing their preferred alleles, male-beneficial alleles that increase mate harm may be constrained from increasing in frequency in the gene pool shared between the sexes. If there is selection for increased harmfulness but the response is constrained by shared genetics, then removing IASC by separating the gene pools of males and females would allow mate harming alleles to increase in frequency among the gene pool experiencing male-limited selection. Previous experimental evolution regimes which separated the gene pools of the sexes (thereby removing IASC) have yielded mixed support for this, with one set of experimental populations showing increased male harm evolving within a gene pool subject to male-limited selection (Rice, 1996; Rice, 1998) but another set of experimental population showing no change (Jiang et al., 2011).

Here we revisit this question with a novel set of experimentally evolved populations, using a design that allowed for a more detailed assessment of male effects on female fitness. We compared levels of mate harm among *Drosophila melanogaster* males that had been subject to 120 generation of sex-limited experimental evolution. For one pool of genetically variable Chromosome 2s, we labelled copies with a fluorescent *DsRed* marker (hereafter “*Red* chromosomes”) and forced these *Red* chromosomes into a Y like inheritance pattern (passed only father to son), experiencing male-limited selection. The alternate pool of non-fluorescent chromosomes (hereafter “*NonRed* chromosomes”) segregated like an X, spending ∼2/3 of their time in females and experiencing female-biased selection. Removing IASC on these *Red* chromosomes, should allow them to increase the frequency of male-beneficial alleles, regardless of their effect when expressed in females. After 120 generations of this sex-limited selection, *Red* and *NonRed* pools showed substantial transcriptional and allelic divergence (Grieshop et al., 2025; Melo-Gavin et al., 2025), and males carrying a *Red* chromosome (hereafter “*Red* males”) displayed increased mating success (Grieshop et al., 2025), suggesting differences in selection between these two pools that had been previously constrained under normal inheritance. If IASC at shared loci has a pleiotropic effect on mate harm under normal inheritance, then we would expect differences in the frequency of mate harm alleles between the male-selected *Red* and female-selected *NonRed* pools due to the relaxation of IASC in this evolution regime.

In addition to investigating the effect of sex-limited experimental evolution on mate harm, we add two further layers of investigation. First, we compare the effect of these male types (*Red* and *NonRed*) on female fecundity across two different levels of male exposure to verify that observed effects on female fecundity are related to mate harm (i.e., that females are more fit when exposure to males is reduced) and are not caused by other effects males may have on females (e.g., differences in benefit to females; see Yun et al., 2021). Further, we examined female fitness following male exposure across three time windows, illuminating unexpected temporal dynamics of mate harm.

We find that females mated to males carrying a male-selected *Red* chromosome had lower fecundity than when mated to males carrying female-selected *NonRed* chromosomes. This reduction in fecundity was lessened when the amount of exposure to males was reduced, indicating differences in male harmfulness. Interestingly, the magnitude and even direction of the effect of male exposure on female fecundity were greatly affected by the time since last male exposure.

## Methods

### Experimental Evolution Protocol

A detailed experimental protocol is given elsewhere (Grieshop et al., 2025). In brief, six replicate populations of *Drosophila melanogaster* (500 males and 500 females per population each generation) were established from a large (*N* = 2000) genetically variable lab population that had been segregating for a dominant fluorescent *DsRed* marker on Chromosome 2 for ∼90 generations prior to the start of the experiment. The marker is located at approximately 2R:48C and no fitness effects of the marker were detected in previous work (Zikovitz and Agrawal, 2013). Hereafter, we refer to copies of Chromosome 2 with and without the DsRed marker as “*Red*” and “*NonRed*” chromosomes, respectively. The males used to sire each subsequent generation were *Red*/*NonRed* heterozygotes and females were *NonRed* homozygotes. That is, *Red* chromosomes were transmitted father to son where, due to a lack of meiotic recombination in males in *D. melanogaster*, they experienced “Y-like” inheritance and male-limited selection. Conversely, *NonRed* chromosomes (spending 2/3 of the time in females) experienced “X-like” inheritance and thereby female-biased selection. A small fraction of each population underwent slightly altered protocols to allow for a low level of recombination on *Red* chromosomes, both between one another and between *Red* and *NonRed* chromosomes. (Because *NonRed* homozygote females are used in the main cross each generation, *NonRed* chromosomes recombine with one another at a higher rate).

We also utilized control populations (not reported in Grieshop et al. (2025)) derived from the same source as the experimental populations. In our six replicate control populations (*N* = 1000 each, 50 males:50 females per bottle with 40mL standard yeast-sugar media), fluorescent flies (*DsRed* marker /+) were paired with nonfluorescent (+/+) flies of the opposite sex. Specifically, each generation 5 bottles contained 50 fluorescent males (*DsRed* marker /+) and 50 nonfluorescent females (+/+) and the other 5 bottles contained 50 fluorescent females (*DsRed* marker /+) and 50 nonfluorescent male (+/+). In this way, both *DsRed* and nonfluorescent chromosomes spent exactly half of their time in each sex. Control populations were otherwise maintained on the same cycle and in the same conditions as the experimental populations (two-week cycle, 50% relative humidity etc.) as reported in Grieshop et al. (2025). These populations were terminated after generation 58.

### Experiment 1: Two Male Type Mate Harm Assay (one exposure level)

At generations 34 and 41 of experimental evolution we assayed mate harm by comparing differences in female fitness when kept with males possessing either a male-selected *Red* chromosome (*Red*/*NonRed* heterozygotes hereafter “*Red* males”) or two female-selected *NonRed* chromosomes (hereafter “*NonRed* males”). This assay was conducted with the same number of replicates in two experimental blocks, one at generation 34 and one at generation 41.

Females that were 2-4 days post-emergence were taken from the ancestral population and continuously exposed to focal males (10F:10M per vial; all males were of a single type, *Red* or *NonRed*) for 3 days, mimicking regular maintenance conditions. There were 30 replicates for each male type per replicate population (split evenly over the two experimental blocks). After the mating phase, 3 randomly chosen females from each replicate vial were transferred into new individual vials to oviposit for 24 hours. Transferring females to individual vials was done to keep density low and minimize any effects of larval competition on survival. Total female fecundity was measured as total adult offspring produced per female by day 12 post oviposition.

Mate harm was also assayed in control populations where the *DsRed* marker did not evolve in a sex-specific manner. In control populations we compared fluorescent (*DsRed* marker/+) and nonfluorescent (+/+) males in the same way as described above.

Female fitness was analyzed separately for control and experimental populations, both using a linear model with Male Genotype, Block (corresponding to experimental evolution generation 34 or 41) and Population as main effects using the *lme4* package in *R* (Bates et al., 2015). The analysis of female survival during the mating phase was performed the same way.

### Experiment 2: 2 Male Types by 2 Exposure Levels Mate Harm Assay

During generations 120 to 125, another mate harm assay was conducted by measuring female fecundity under both normal (continuous) and reduced (periodic) exposure to males carrying either a male-selected *Red* or two female-selected *NonRed* chromosomes. If males are harmful to females, females should have higher fitness when they experience reduced exposure to males.

Three-day post-emergence virgin females from the ancestral population were placed with focal male flies (10M: 10F per vial, all males were of a single type) for 3 days mimicking maintenance conditions. In the continuous (normal) exposure treatment females were continuously exposed to males. In the reduced exposure treatment, two 4-hour blocks of male exposure occurred, one at the beginning and one at the end of the 3-day mating period. For this treatment, males were removed between exposure episodes and were maintained on fresh food. The same pool of males was used for both exposure episodes. In both exposure treatments, females and males were handled similarly so that they received the same amount of CO_2_ over the course of the mating phase (i.e., continuous exposure replicates where sham treated to mimic the handling and CO_2_ required for the reduced exposure treatment). There were ∼30-35 replicates for each male type for each exposure level per replicate population.

After exposure to males, 8 randomly chosen females per replicate vial were transferred to oviposition vials and allowed to lay eggs for a total of 48 hours, broken up into three time windows (transferring to a new vial for each time window): 0-20 hours after last male exposure (hereafter, “first period”), 20-24 hours after last male exposure (“middle period”), and 24-48 hours after last male exposure (“end period”). The same 8 females stayed together across the entire oviposition window. However, due to deaths during the mating phase, initial oviposition vials for 6 replicates in the continuous exposure condition from populations 4 (3 containing *Red* males and 3 containing *NonRed* males) were supplemented with 1 additional female from neighbouring vials to make 8.

Total female fecundity across the 48-hour oviposition period was measured as total adult offspring per ‘expected’ female produced by day 15 after oviposition. While oviposition vials all start with 8 females in the first period, some females escaped their vials due to experimental error between vial transfer such that some oviposition vials in later time periods had 7 instead of 8 flies. As such total fecundity per ‘expected’ female (i.e., excluding females lost by experimental error) was calculated separately for each time period and then added together to account for differences in number of laying females. If a female died after being flipped into a vial (that is, by some cause other than experimental error) it was still counted as a laying female.

For comparisons across the three time periods, fecundity was standardized as total offspring produced per female per hour to account for differences in duration of time windows. Total fecundity was analyzed with the *lme4* package in *R* using a linear mixed model (Bates et al., 2015) and data across time periods was analyzed using a Repeated Measures Anova using the *rstatix* package in *R* (Kassambara, 2023). Male Genotype, Exposure Level, Population and Time Window (when appropriate) were treated as main effects. Replicate populations were assayed in different experimental blocks (one population per generation over generations 120-125) so variation associated with “Population” represents both genetic and any block differences among populations. However, our main interest is in effects within populations (e.g., male type, exposure level).

## RESULTS

While our selection regime removes IASC for male-selected *Red* chromosomes, selection for mate harm itself is not directly manipulated in our design. If IASC has no effect on mate harm, then female fitness should be similar regardless of mating partner. Alternatively, if IASC has a pleiotropic effect on mate harming alleles and acts to constrain their evolution, then removing IASC in the male-selected *Red* chromosome pool will allow mate harming alleles to increase in frequency, leading to lower fitness when females mate with *Red* males compared to *NonRed* males.

In Experiment 1, at Generation 34 and 41, ancestral females exposed to male-selected *Red* males had significantly lower fecundity than when exposed to female-selected *NonRed* males, suggesting male-selected chromosomes were more harmful to female fitness (*F*_1,348_ = 4.231; *p* = 0.0398; **FIGURE 1**). When using males from control populations in which the *DsRed* marker had not evolved in a sex-specific manner, females performed similarly when exposed to fluorescent (*DsRed* marker/+) and nonfluorescent (+/+) males, implying that the difference between *Red* and *NonRed* male types observed in experimental populations was not due to possession of the fluorescent *DsRed* marker. Female survival during the mating phase was not significantly affected by the male type they were mated to suggesting the mate harm we observed operated primarily on fecundity (**SUPPLEMENTAL FIGURE 1**). The reduced fitness of females exposed to *Red* rather than *NonRed* males is consistent with *Red* males being more harmful. However, a second alternative is that the *NonRed* males were beneficial to females rather than *Red* males having been more harmful. Assays like the one above comparing female fitness when exposed to different male types cannot distinguish between differences in harm vs. differences in benefits (Yun et al., 2021).

**FIGURE 1:**
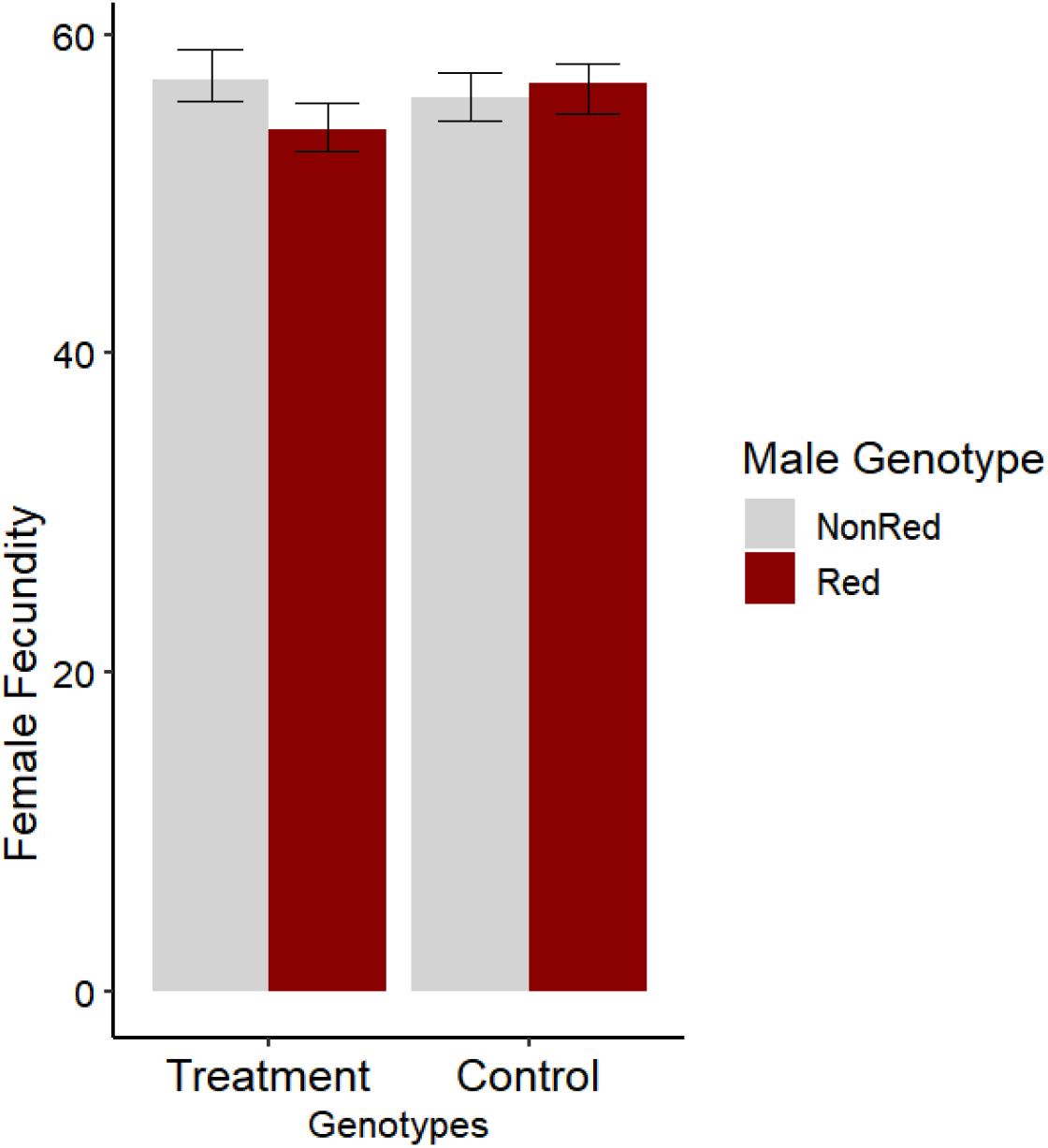
Mate Harm in Experiment 1. Differences in female fecundity when mated to males carrying a male-selected *Red* chromosome or two female-selected *NonRed* chromosomes (Treatment group) compared to control. Grey bars represent *NonRed* males and red bars represent *Red* males. Error bars represent the 95% confidence interval of the mean.

To address this, at generations 120-125, an additional experiment (Experiment 2) was conducted by measuring ancestral female fecundity under both reduced (periodic) and normal (continuous) exposure to both male genotypes. Effects of mate harm should decrease with reduced male exposure, whereas genetic effects affecting offspring viability should remain constant across exposure levels. Moreover, in this experiment, we partitioned female fitness across three periods spanning 48-hours following the male exposure manipulation. This allowed us to examine the effects of male exposure and the differences between the effects of the male-selected *Red* and female-selected *NonRed* chromosome pools in more detail.

When examining total female fecundity over the 48-hour oviposition window, reduced exposure to males was associated with significantly increased fecundity, both when averaged across male types (*F*_1,811_ = 4.39; *p* < 0.05; **FIGURE 2A;** see **Table 1** for full model) or examined individually (*NonRed*: *F*_1,405_ = 4.35; *p* < 0.05; *Red*: *F*_1,405_ = 36.30; *p* < 0.0001), indicating both male types were harmful (verifying that the difference between the effect of exposure to *NonRed* rather than *Red* males in first experiment was not because *NonRed* males were “beneficial” to females). The effect of this mate harm varied significantly by genotype (Exposure by Genotype: *F*_1,811_ = 7.61; *p* < 0.001), where females mated to males carrying a male-selected *Red* chromosome had lower fecundity at continuous exposure than when mated to males carrying female-selected *NonRed* chromosomes. Examining female fecundity only at continuous exposure, we recapitulated the significant genotype effect from Experiment 1 (*F*_1,397_ = 5.92; *p* < 0.05). Collectively, these results indicate *Red* males were more harmful than *NonRed* males.

**Table 1:**
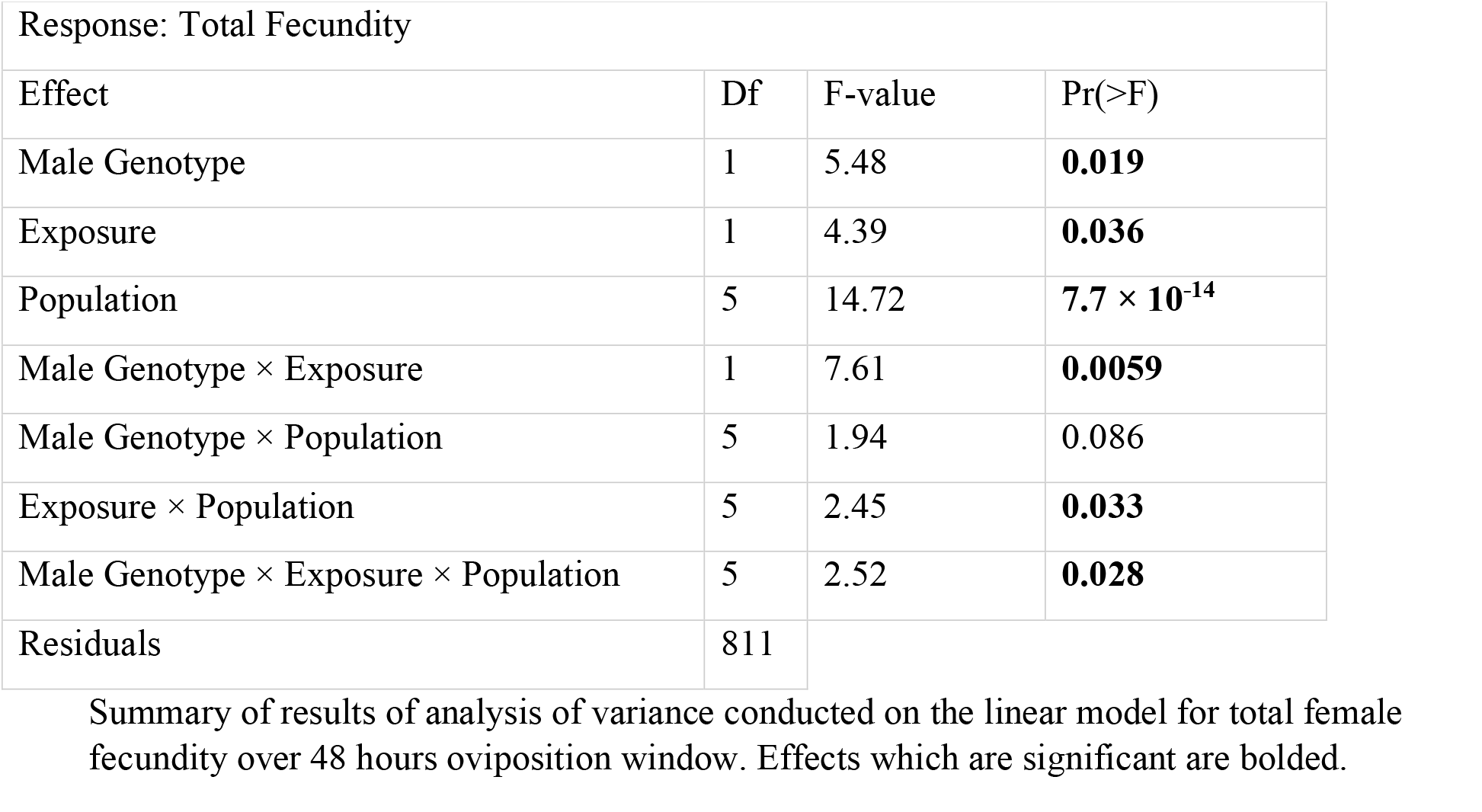
Results for the Analysis Total Fecundity for Experiment 2.

**FIGURE 2:**
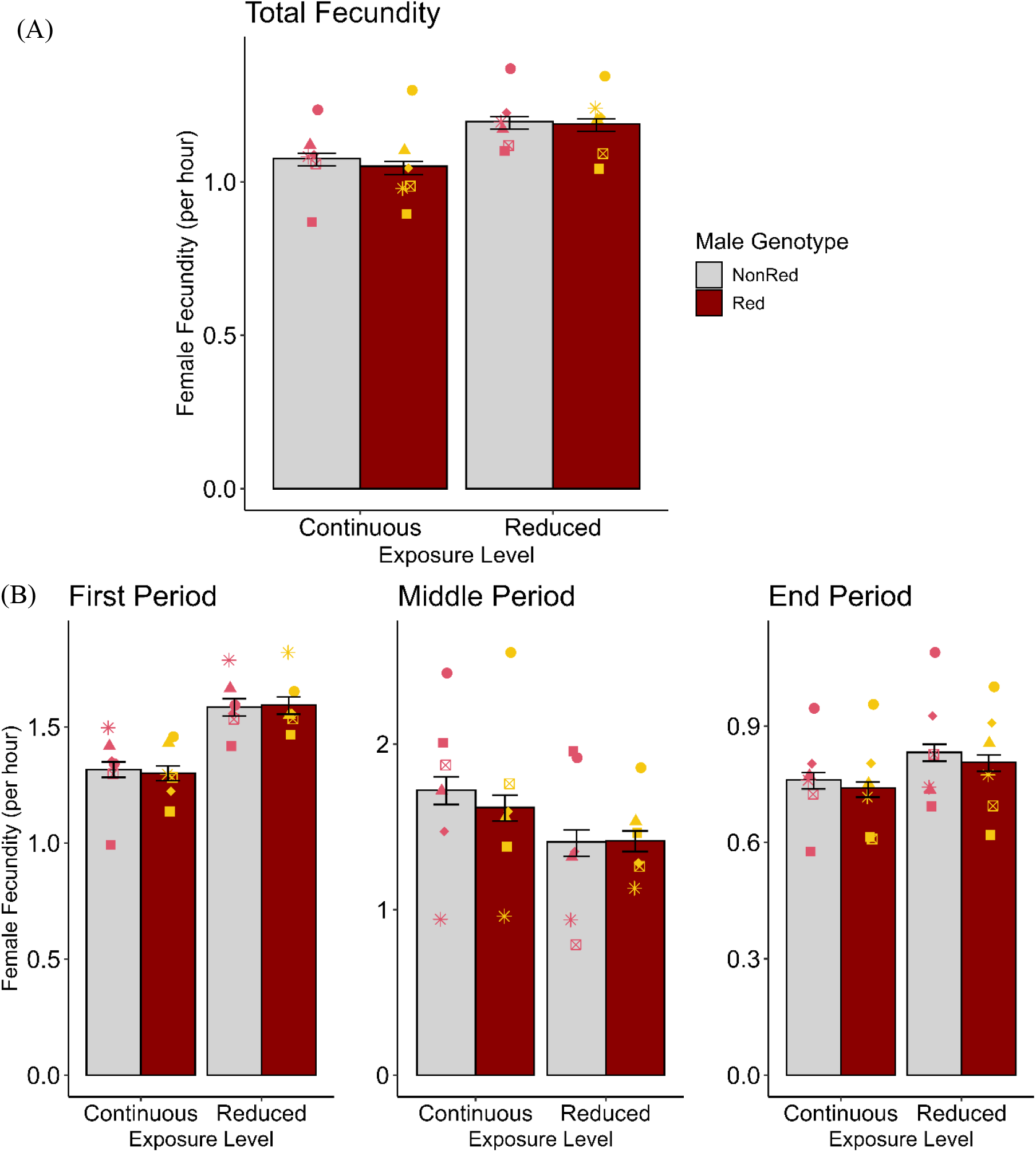
Mate Harm in Experiment 2. Differences in female fecundity when mated to males carrying either a male-selected *Red* chromosome or two female-selected *NonRed* chromosomes, at either normal (continuous) or reduced (periodic) male exposure. (A) Total Fecundity of females for the entire 48-hour oviposition period. (B) Female Fecundity examined over time windows. (First period = 0-20h, middle period = 20-24h, end period = 24-48h). Yellow and pink points represent replicate population means. Error bars represented bootstrap 95% confidence interval of the mean.

However, it should be noted that this effect varied significantly among populations (Genotype by Exposure by Population: *p* < 0.05; see **Table 1** for full model). Though only one population showed the opposite trend to that described above (**FIGURE 2A**), the replicate populations were highly variable in terms of fecundity (averaged across male types and exposure levels; Population Effect: *F*_5,811_ = 14.72; *p* < 0.001) and the strength of the effect of exposure (Population by Exposure: *F*_5,811_= 2.45; *p* < 0.05). As population and experimental block were conflated in this experiment for logistical reasons, we cannot determine whether the “population” effects reflect true genetic differences among replicate populations.

Examining female fecundity across the time windows (**FIGURE 2B**) revealed the effect of exposure was highly sensitive to the time of measurement (*F*_2,2430_ = 813.1; *p* < 0.001; **FIGURE 2B; TABLE 2** for full model). There was a significant effect of exposure in the first period in the expected direction (that is, indicating mate harm) (*F*_1,811_ = 13.58; *p* < 0.001). However in the shorter middle period, this effect reversed such that females continuously exposed to males had higher fecundity than those with reduced male exposure (though for half of the populations this “reversal” only occurred for females mated to *NonRed* males) (Exposure by Population effect: *F*_5,811_ =7.40; *p* < 0.001 and Exposure by Genotype by Population effect: *F*_5,811_ = 2.43; *p* < 0.05). During this “reversed” middle period, *NonRed* males were more beneficial than *Red* males (Genotype by Exposure: *F*_1,811_ = 3.81; *p* = 0.051; and Genotype by Exposure by Population: *F*_5,811_ = 2.43; *p* < 0.05). By the end period, the effect of exposure was once again in the expected direction, though dampened on average, and varying in strength among populations (Exposure by Population effect: *F*_5,811_ = 4.45; *p* < 0.001).

**Table 2:**
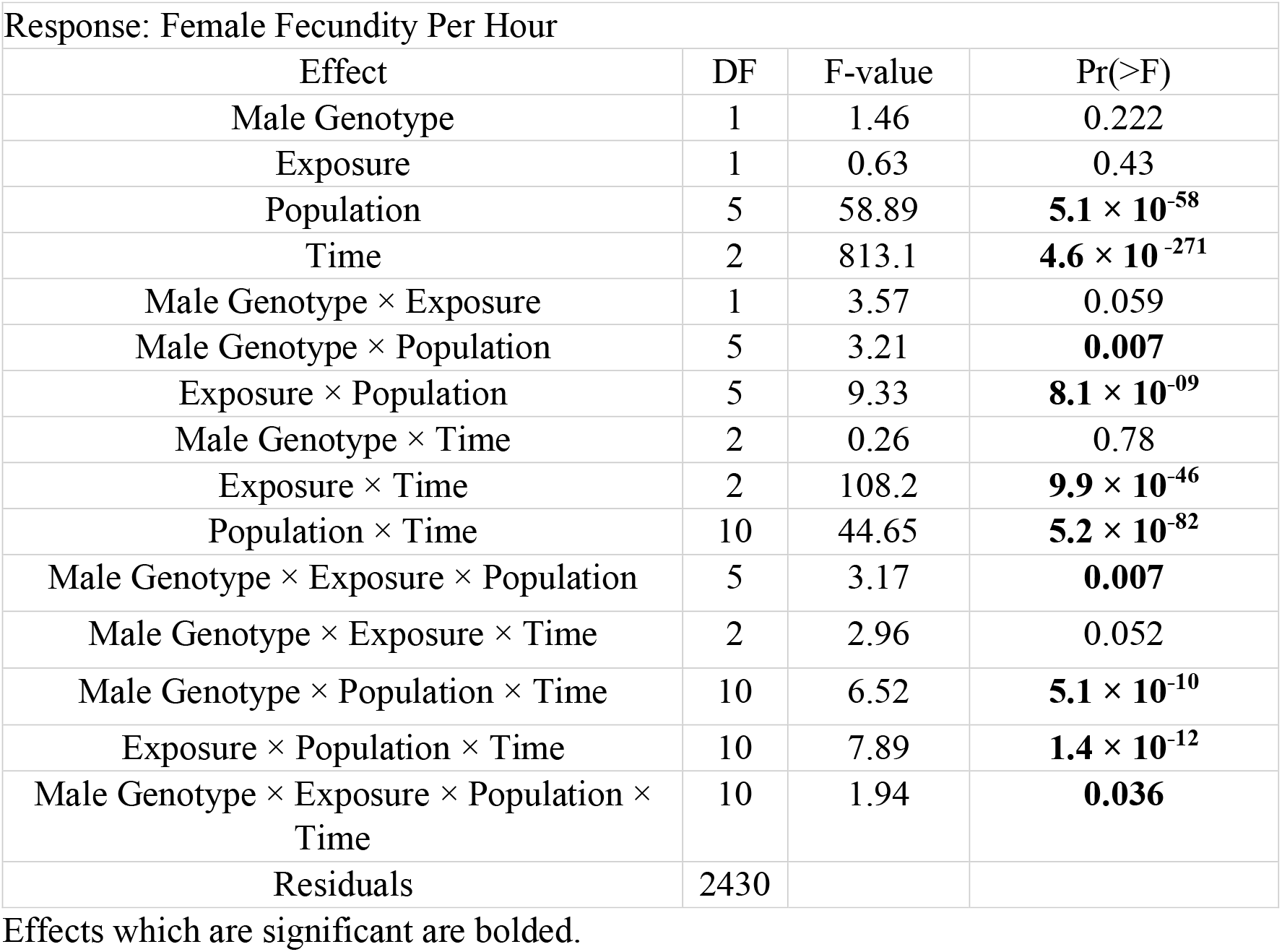
Experiment 2 Results for Repeated Measured Analysis of Variance (ANOVA) of Fecundity Per Hour Across 3 Time Windows.

Lastly, we also observe a significant four-way interaction between Genotype, Exposure Level, Population and Time (*F*_10,2430_ = 2.941; *p* < 0.05). This is not surprising given the three-way interaction we observed when examining the middle period individually, and how different the middle period is from the other two.

## Discussion

In many species, competition occurs among males to fertilize females. This generates different selection on males than on females. At the genetic level, this can result in alternative alleles being favoured in each sex (IASC). In some cases, at the phenotypic level, it may result in males evolving traits that are harmful to the females with whom they interact. Yet, little empirical work has focused on the interaction of IASC and mate harming traits. If there is selection for increased mate harm, but mate harming alleles are constrained by IASC, then separating the gene pools of the sexes by imposing male-limited selection would allow these alleles to increase in frequency and cause males to be more harmful.

Here we imposed sex-biased selection creating two pools of genetically variable chromosomes, one male-selected *Red* chromosome pool and one female-selected *NonRed* chromosome pool. After 120 generations of experimental evolution, these pools were diverged with respect to gene expression, allele frequency, and male mating success (Grieshop et al., 2025; Melo-Gavin et al., 2025), suggesting adaptation to sex differential selection occurred in these pools that had been prevented under normal inheritance. If loci underlying mate harming traits are affected by IASC, our relaxation of this conflict should lead to differences in mate harming ability between males carrying male-selected *Red* chromosomes or female-selected *NonRed* chromosomes.

Mating with *Red* males reduced the fecundity of females more than mating with *NonRed* males suggesting that interacting with *Red* males was more harmful to females. Though we cannot know if *Red* males evolved to be more harmful or *NonRed* males evolved to be less harmful, either change would indicate a pleiotropic effect of IASC on mate harm. We suspect the most likely cause of this result is that there is some degree of opposing selection on alleles affecting mate harm between the *Red* and *NonRed* pools. An alternative is that divergence in the *NonRed* pool occurred via the relaxation of purifying selection on variants with male effects but no fitness effects when carried by females. This possibility seems unlikely given *NonRed* chromosomes spent one third of their time in males, exposing them to some male selection.

Though we detected an average effect of increased harm in *Red* compared to *NonRed* males, there was considerable variation among populations. As different populations were assayed in different experimental blocks, we cannot determine if the among-population variation reflects genetic differences or is due to subtle (unknown) differences in assay conditions. Nonetheless, this variation may suggest that, while there tends to be a relationship between mate harm and IASC, this relationship can be absent, weak, or context dependent.

Perhaps the variation we observed among populations echoes the mixed results from previous experiments employing sex-limited selection. Using the same ancestral population as each other, Rice (1996) and Prasad et al. (2007) both performed experimental evolution to create a pool of male-limited haplotypes, though using a different method from our study. While both groups found males possessing a male-selected haplotype had increased fitness compared to controls (Prasad et al., 2007; Rice, 1996; Rice, 1998), the effects of these male-selected haplotypes on female fitness were mixed. Rice (1996, 1998) found that when males carrying male-selected haplotypes mated with females (tester females carrying ancestral LH_M_ background (Rice, 1998)), female mortality was significantly higher than when females mated with control males. Jiang et al. (2011) using the populations evolved by Prasad et al. (2007) and the same assay design as Rice (1998) found no difference in female mortality when exposed to males with male-selected haplotypes compared to control haplotypes. Collectively, the existing empirical evidence suggests that separating the gene pool to allow sex-specific responses to selection tends to affect mate harm on average, though this may not be a deterministic process and other context dependent factors may play a large role in the evolution of mate harming ability.

A common approach in studying mate harm is to compare female fecundity when mated to different male types under continuous exposure. In this approach, if females have lower fecundity with male type A, that male type is interpreted as the more harmful one. However, differences in the effect of males on females may occur for two additional classes of reasons other than mate harm. First, there may be differences in “genetic quality” (i.e., differences in male (in)fertility or compatibility) that may impact female fecundity. For example, an alternative interpretation of the result that females have higher fitness with *NonRed* males is that *Red* males are more likely to be infertile or genetically incompatible or have offspring less likely to survive. A second alternative is that the difference arises because male types differ in benefit to their mates rather than a difference in harm. Though *Drosophila melanogaster* is often viewed as a classic example of male harm, males affect females in a variety of ways, not all of which are deleterious (Ram and Wolfner, 2007a). The net effect of males on females can be dependent on context, including the environment in which interactions occur and the diet and life history of the individuals (Yun et al., 2017; Londoño-Nieto et al., 2023; Fricke et al., 2010; Yun et al., 2021).

The inclusion of an exposure level manipulation in Experiment 2 helps address these alternatives. The exposure manipulation provided a clear signal that both *Red* and *NonRed* males were net harmful rather than beneficial (i.e., total female fecundity was lower when females had continuous rather than reduced exposure to males, regardless of male type). Notably, differences between the effect of *Red* versus *NonRed* males on female fecundity were only significant under continuous male exposure, not under reduced exposure, suggesting these differences were driven by male interaction with females. (In contrast, a difference in genetic quality of male types should result in similar fitness differences at both exposure levels, rather than the significant Genotype by Exposure effect we observe.)

We found that the effect of male exposure on female fecundity was highly sensitive, in magnitude and even direction, to the time since last mating. While we do not know the cause of these changes over time, several post-copulatory traits known to affect mate harm have time sensitive effects that may initially increase egg laying but reduce later or lifetime fecundity (Ram and Wolfner, 2007a; Ram and Wolfner, 2007b; Chapman et al., 1995). In this vein, the reversal of the sign of mate harm between 20-24 hours since last male exposure could be consistent with effects of sperm proteins temporarily increasing fecundity before leading to harm later. Indeed, many seminal fluid proteins have effects peaking at 24 hours (Sirot et al., 2014; Heifetz et al., 2001). Given that male-limited selection can produce changes in sperm competition (Rice, 1998) and that the outcome of sperm competition depends heavily on the genotypes of the competing males as well as the genotypes of females (Clark and Begun, 1998; Clark et al., 1999), it is conceivable that there population-specific differences in post-copulatory traits that are responsible for the Genotype by Exposure by Time by Population effect observed here.

Overall, females mated to males carrying male-selected *Red* chromosomes exhibit lower fecundity than when mated to males carrying female-selected *NonRed* chromosomes. To the extent that males are harmful (that increased exposure decreases fecundity), *Red* males appear to be more harmful, and to the extent that males are beneficial (i.e., during the middle period), *Red* males are less beneficial. This suggests that alleles affecting mate harming ability have changed in frequency in response to our sex-biased selection regime, implicating the role of IASC in constraining the evolution of mate harm. It is becoming apparent that selection on one sex can have profound effects on the other sex, even with respect to sex-limited traits. Understanding the role and underlying architecture of shared genetics may be necessary in understanding the co-evolutionary dynamics of mate harm (Pennell et al., 2016).

Many species exhibit mate harm (e.g., elephant seals (Le Boeuf and Mesnick, 1991), water striders (Arnqvist and Rowe, 2002), cowpea weevils (Gay et al., 2010)), and though the underlying genetic architecture of these mate harming traits is often not well characterized, there is ample reason to believe IASC may play a role in modulating mate harm. In the cowpea weevil, a quantitative genetic analysis indicated that alleles that increased a male’s harmfulness, also decreased longevity when expressed in females, suggesting a mechanism by which selection for increased mate harm could be constrained by deleterious fitness effects when in females (i.e., IASC; Gay et al., 2010; see Friberg et al. (2005) for a similar study in *D. melanogaster*). In species such as northern elephant seals, southern elephant seals, and monk seals, males can be three to seven times larger than females and, because of their much larger size, can injure and even kill females during aggressive mating attempts (Le Boeuf and Mesnick, 1991; Deutsch et al., 1990), or in pre-mating abduction attempts (Campagna et al., 1988). This dimorphism in size is driven by intense selection for male-male competition (Haley et al., 1994). Optimal body size for females is presumably smaller than for males, perhaps due to their different foraging strategies (Kienle et al., 2021). Given that there tends to be strong positive genetic correlations in body size in animals (Poissant et al., 2009), this would result in IASC on alleles affecting body size, where these same alleles would also affect mate harm. Understanding the evolution of sex specific adaptations (such as mate harm) also requires understanding the factors that may constrain it, such as shared genetics.

## Acknowledgements

We are grateful for everyone who helped us maintain these populations.

## SUPPLEMENTAL FIGURES

**SUPPLEMENTAL FIGURE 1:**
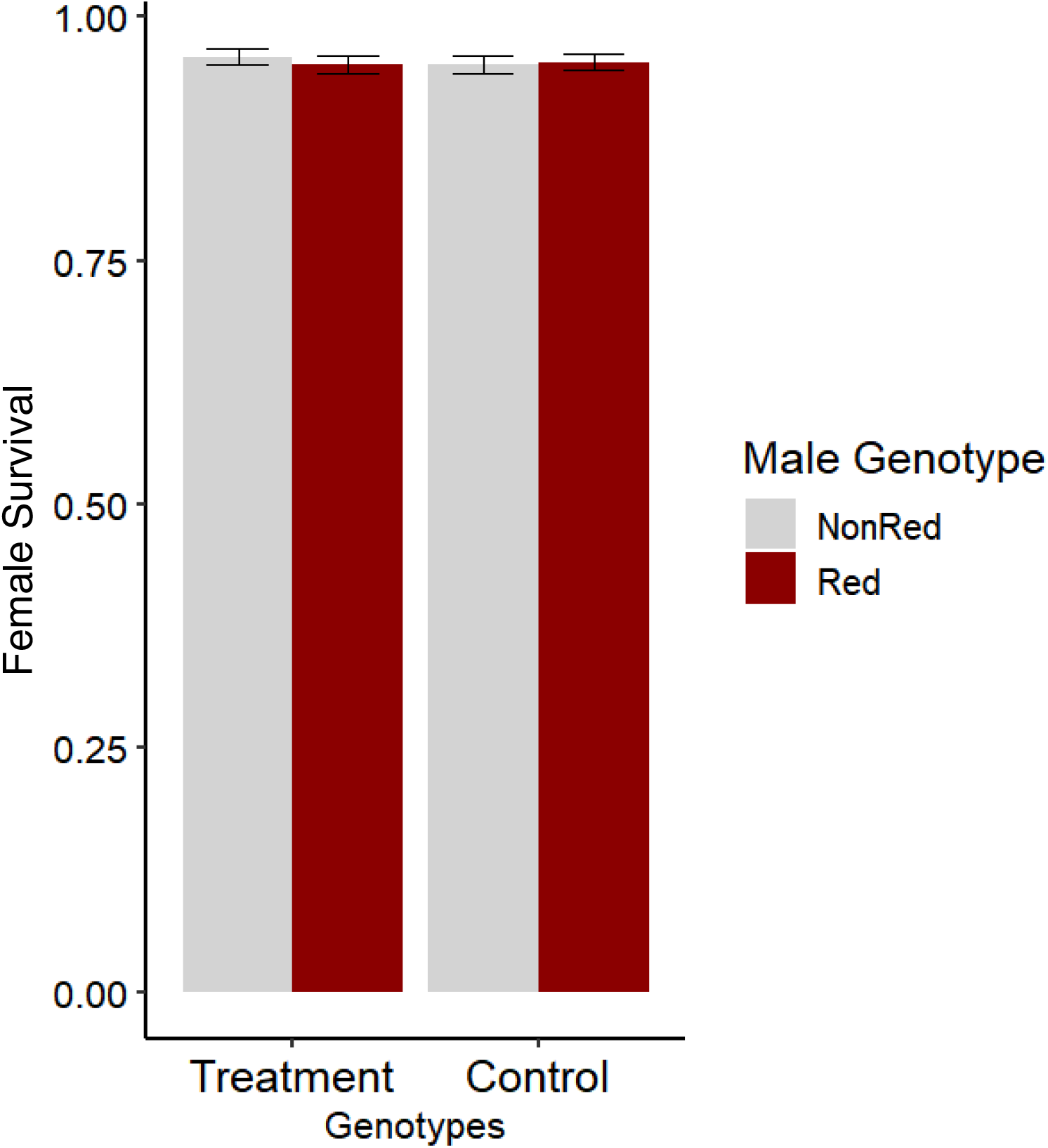
Female Survival after 3 days of mating in Experiment 1. Compares mean female survival after mating with *Red* and *NonRed* genotypes from either experimental (Treatment) or control populations. Errors bars represent ± standard error.

