## Supplemental Figure 1 for "Response of mate harm to sex-separated gene pools"

SUPPLEMENTAL FIGURES

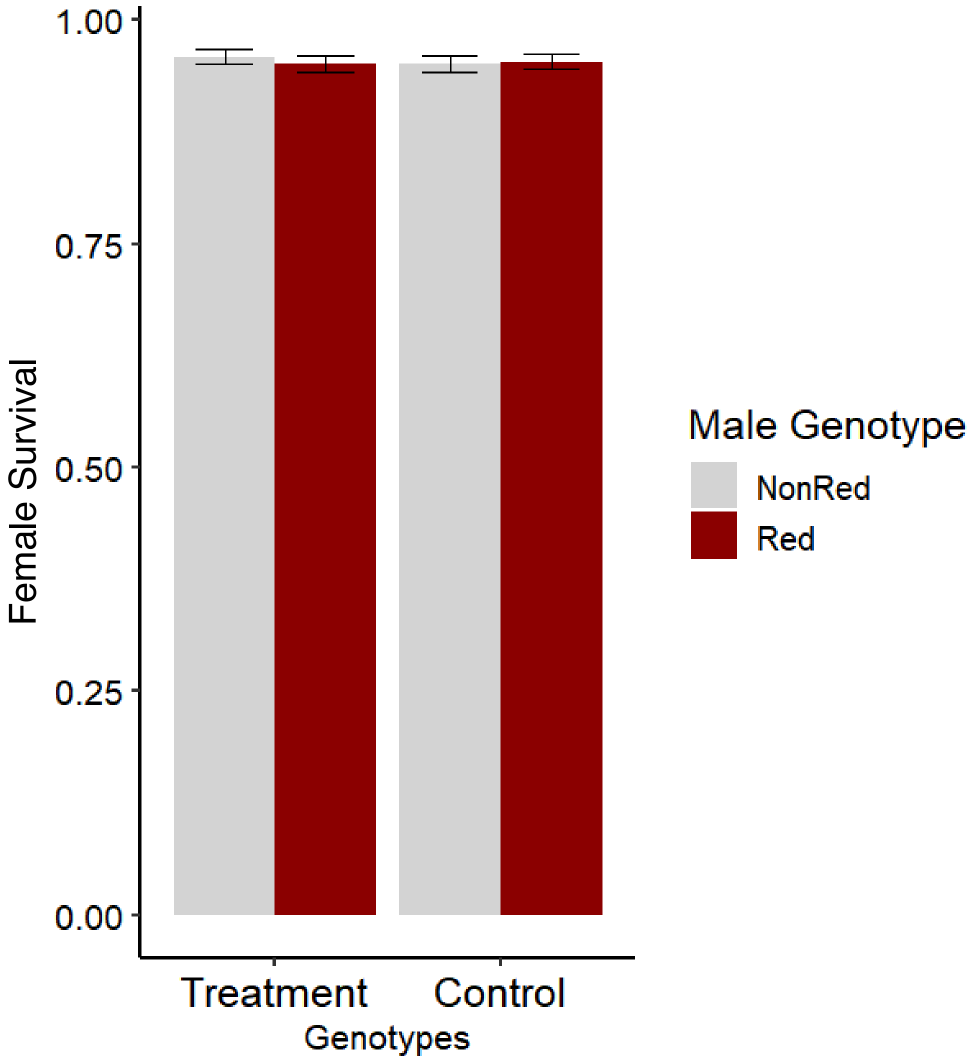


**SUPPLEMENTAL FIGURE 1:** Female Survival after 3 days of mating in Experiment 1. Compares mean female survival after mating with *Red* and *NonRed* genotypes from either experimental (Treatment) or control populations. Errors bars represent ± standard error.
